# A capsule polysaccharide synthesis locus database for the *Klebsiella oxytoca* Species Complex

**DOI:** 10.64898/2026.07.16.739023

**Authors:** Melinda M. Ashcroft, Naoise McGarry, Thomas D. Stanton, Lesley Hoyles, Kathryn E. Holt, Kelly L. Wyres

## Abstract

The *Klebsiella oxytoca* Species Complex (SC) represents an emerging healthcare-associated group of opportunistic pathogens. The capsular polysaccharide is a virulence determinant and target for novel vaccines, monoclonal antibodies and phage therapy. In the absence of broadly accessible phenotyping techniques, prediction of capsule types from whole-genome sequence data is critical for understanding capsule diversity and epidemiology, and to prioritise capsule types as targets for novel anti-*K. oxytoca* SC interventions. Here we present the first comprehensive capsule synthesis locus (K locus) database targeted for the *K. oxytoca* SC, comprising 88 distinct loci defined by gene content and which is compatible with the rapid genome typing tool Kaptive. The database provides high coverage of publicly available *K. oxytoca* SC genomes (97.6% of 2,244 genomes, dereplicated from a total of 4,055), and the typing rate is significantly higher than that achieved with the pre-existing *Klebsiella* K locus database (97.6% vs 50.3%, *p* <0.0001), which primarily targets the *Klebsiella pneumoniae* SC. We demonstrate the utility of the novel *K. oxytoca* SC database by application to three diverse clinical *K. oxytoca* SC isolate collections (n=61 to 102 genomes each), suggesting a high diversity of K types. The novel *K. oxytoca* SC K locus database (github.com/klebgenomics/KoSC-surface-antigen-loci) will provide a key resource to support larger systematic studies and ongoing genomics surveillance efforts for the *K. oxytoca* SC.

**IMPACT STATEMENT:** Members of the *Klebsiella oxytoca* Species Complex (SC) are an emerging cause of infections in humans and are frequently associated with antimicrobial resistance. *Klebsiella* species produce two key surface antigen sugars (capsular polysaccharide and lipopolysaccharide) that are immunogenic and are targets for novel control strategies such as vaccines and phage therapy. Phenotypic typing of these surface antigen sugars (serotyping) is costly and laborious, with genotyping (predicting the serotype from whole-genome sequence data) a useful alternative. Here, we present a curated capsule (K) locus reference database for the *K. oxytoca* SC, which represents a useful tool to assist in the epidemiological surveillance of this emerging pathogen.

**DATA SUMMARY:** All *Klebsiella oxytoca* Species Complex genomes used in this work were publicly available, with accession details listed in **Supplementary Tables 1, 2 and 3**. The *K. oxytoca* Species Complex K locus reference database is available under GNU public license at github.com/klebgenomics/KoSC-surface-antigen-loci.

## INTRODUCTION

Members of the *Klebsiella oxytoca* Species Complex (SC) are commensals of human-associated microbiotas, from where they can emerge as opportunistic pathogens causing antibiotic-associated haemorrhagic colitis, skin, wound and soft-tissue infections, pneumonia, urinary tract and systemic infections [1]. The complex includes *K. oxytoca, Klebsiella michiganensis Klebsiella grimontii* and *Klebsiella pasteurii*, which appear to be the most clinically relevant [1–3], as well as *Klebsiella spallanzanii, Klebsiella huaxiensis* and three unnamed taxa [4]. *K. oxytoca* SC outbreaks and nosocomial infections are becoming a public health problem [5], with reports of increasing antimicrobial resistance (AMR). Notably, carbapenem and third-generation cephalosporin-resistant strains are among those designated as urgent priority pathogens by the World Health Organization because they cause infections that are extremely difficult to treat [6].

Multi-valent vaccines, monoclonal antibodies, and phage therapies targeting the *K. oxytoca* SC polysaccharide capsule represent alternate strategies to combat AMR. In *Klebsiella* spp., the capsular polysaccharide (K antigen) is a key pathogenicity factor and useful epidemiological marker [7, 8]. The capsule protects *Klebsiella* spp. from phagocytosis and biocidal molecules and provides resistance to complement-mediated killing [9–12]. Multi-valent protein-conjugate polysaccharide vaccines have been shown to be highly effective against *Streptococcus pneumoniae* and *Neisseria meningitidis* [13, 14]. Notably, capsule polysaccharide vaccines against *K. pneumoniae* – a close relative of the *K. oxytoca* SC and the most common cause of *Klebsiella* spp. infections – were previously shown to be protective in mice and humans [15, 16], and several such vaccines are currently in development [17]. Similarly, anti-*K. pneumoniae* monoclonal antibody therapies targeting the capsule were recently shown to be effective in a mouse model [18].

In *K. pneumoniae* and closely related organisms in the *K. pneumoniae* SC, there is considerable antigenic diversity of the capsule polysaccharide, with 82 serologically defined capsule types [19–22]. The original *Klebsiella* serotype reference strains are often considered as *K. pneumoniae*, but six are now known to represent members of the *K. oxytoca* SC. Five of these six *K. oxytoca* SC have known capsule polysaccharide structures (K26 [23], K41 [24], K66 [25], K70 [26], K74 [27]). A novel structure and corresponding novel capsule (K) locus were also recently described for *K. grimontii* strain K15g [28], but to our knowledge no other *K. oxytoca* SC capsular polysaccharide structures have been described.

Whole-genome sequencing enables high-resolution characterisation, genotyping and surveillance of bacterial pathogens, and is an established approach for predicting capsule types based on the K locus [22, 29, 30]. In the *K. pneumoniae* SC, the K locus is typically flanked by *galF* and *ugd* genes, 16–25 kb in length and comprising 18–25 open reading frames (ORFs), with an average GC content of 40%, which is substantially lower than the surrounding chromosome (average GC content of 55–60%) [31, 32]. The general arrangement of the K locus is a terminal region and a central region. In the terminal regions is a set of conserved genes that form the core capsule biosynthesis machinery (e.g., *galF, wzi, wza, wzb, wzc, gnd* and *ugd*) [33]. The central region carries a set of accessory genes that are highly variable, encoding sugar synthesis, biochemical modifications of these sugars, processing (glycosyltransferases) and export proteins, plus the core assembly components *wzx* (encoding a flippase) and *wzy* (encoding a capsule repeat unit polymerase), of which there are numerous variants [32, 33]. Genomic analyses have revealed >162 distinct *K. pneumoniae* SC K loci defined on the basis of unique gene content within the central region of the locus [32, 34], each of which is predicted to result in the production of a distinct polysaccharide structure.

Our team has previously described a standardised nomenclature for *K. pneumoniae* SC K loci and developed Kaptive, a bioinformatics tool that rapidly characterises these loci from whole-genome sequences, taking as input a curated reference database of locus sequences, and query genome assemblies [30, 32, 34]. The database was labelled as the *Klebsiella* K locus database because it captured loci from *K. pneumoniae sensu stricto* as well as the other taxa within the *K. pneumoniae* SC (plus the subset of the original *Klebsiella* serotype references that are now known to represent more distantly related *Klebsiella* species). The first version of the database also included two loci from *K. oxytoca* genomes and one from an unclassified *Klebsiella* spp. genome (derived from a total of 10 *K. oxytoca* and four unclassified *Klebsiella* spp. genomes that were available at the time) [32]; however, subsequent updates have focussed solely on *K. pneumoniae* SC, as the primary focus of the database [34–36]. Other teams have reported the application of the database to *K. oxytoca* SC assemblies [37–40] and reported low typeability rates of 0-33%. It is, therefore, clear that the original database – hereafter called the *K. pneumoniae* SC database – does not provide adequate coverage of the *K. oxytoca* SC population, and we caution against its use in this context because Kaptive is optimised for databases with high population coverage. Notably, genomes harbouring loci that are not represented in the reference database can be mistyped when using Kaptive’s default settings [30].

To address the lack of a suitable database for typing *K. oxytoca* SC K loci, we collated publicly available *K. oxytoca* SC genomes and extracted and curated all contiguous K loci. We assigned a standardised nomenclature and generated a Kaptive-compatible reference database, for which our analyses confirm high coverage of available *K. oxytoca* SC genomes. We further demonstrate the utility of this database to support *K. oxytoca* SC seroepidemiology analyses by application to three clinical isolate genome collections.

## METHODS

### Genome sequences and quality control

We generated a preliminary dataset at the initiation of this study in 2022 (comprising short- and long-read data, most of which we assembled *de novo*, see below), from which we identified a preliminary set of *K. oxytoca* SC K loci. During the course of this work, numerous additional *K. oxytoca* SC genomic sequences were deposited in public archives and novel standardised short-read assembly collections were created by third parties [41]. Furthermore, the taxonomic relationships within the *K. oxytoca* SC were further clarified and novel taxa identified, including “*Klebsiella mammalorium*” (carrying the *bla*^OXY12^ beta-lactamase gene [4], formal description in progress, LH and KLW) [42]. We therefore sought to leverage these standardised assembly collections to expand the *K. oxytoca* SC K locus database and generate K locus typing information that will be of benefit to other teams using the standardised genome assembly collections. We used the updated dataset for all subsequent analyses and refer to it hereafter as the final dataset. For transparency, we describe below the assembly and quality control procedures for both datasets.

### Preliminary dataset

We downloaded 1,667 *K. oxytoca* SC genome assemblies that were publicly available from Pathogenwatch (https://pathogen.watch/, n=598) and the NCBI non-redundant (n=733) and whole-genome sequence databases (n=336; https://www.ncbi.nlm.nih.gov) as of 26 October 2022. We also obtained short-read archive run metadata for all *K. oxytoca* SC accessions. We excluded duplicates and long-read only sequence data and downloaded the corresponding raw sequence reads of 1,092 isolates.

Additionally, 776 genome assemblies were collated from published studies [2, 3, 43]. Raw Illumina sequence reads were trimmed using Trim Galore v0.5.0 (https://github.com/FelixKrueger/TrimGalore, leveraging cutadapt v2.7 [44]) with default parameters, excluding reads that had a mean Phred quality score ≤20. For isolates with only Illumina sequencing data, trimmed reads were assembled using Unicycler v0.4.7 [45] with default parameters. For isolates with paired long-read (Pacific Biosciences (PacBio) or Oxford Nanopore Technologies (ONT)) and Illumina short-read sequencing data, the long reads were assembled *de novo* using Flye v2.8 [46], using a genome size of 6 Mb and the “--asm-coverage 40” flag to subset the longest reads for initial disjointing assembly, with five polishing iterations and otherwise default parameters. Long-read assemblies were further polished by mapping the corresponding, trimmed Illumina reads to each contig using bwa mem v0.7.17 [47] and samtools v1.9 [48] and then correcting for single nucleotide polymorphisms (SNPs) and insertions and deletions (indels) with Pilon v1.24 [49] until no changes remained.

All genome assemblies were then assessed using Kleborate v2.2.0 [50] with default parameters and were excluded from the study if the genomes were non-*K. oxytoca* SC species, or if the assemblies failed the empirically determined quality control criteria: (i) total assembly length <4.5 Mb or >7.5 Mb; (ii) contig number >500; (iii) N50 statistic <20 Kb; (iv) the number of ambiguous bases (non ACGT) >50 Kb; (v) evidence of >1% non-*Klebsiella* read contamination as determined by MetaPhlAn v4.0.6. Where a BioSample contained multiple SRA accessions of the same sequencing platform, assemblies were compared based on QC metrics, with only the best assembly kept. A total of 3,288 genome assemblies remained after QC filtering for use as the preliminary dataset.

Accession and QC details are available in **Supplementary Table 1**.

### Final dataset

We retrieved genome assemblies from NCBI (https://www.ncbi.nlm.nih.gov), as well as the Pathogenwatch (https://pathogen.watch) and AllTheBacteria [41] collections as of 5 April 2025. The latter two collections comprise standardised assemblies of read data drawn from the European Nucleotide Archive.

Assemblies were excluded based on the species-specific thresholds defined by the newly developed Qualibact approach where available (https://qualibact.org/Klebsiella, all except “*K. mammalorium*” and the unnamed taxa), or otherwise empirically determined quality control criteria (genome size >7 Mbp, and >800 contigs). Assemblies were then dereplicated by Biosample ID, resulting in a final dataset of 4,055 whole-genome assemblies. Notably, this updated dataset included representatives of all taxa currently considered part of the *K. oxytoca* SC as well as the closely related species *Klebsiella indica* (**Supplementary Figure 1**). Isolate metadata, including sample source, country of origin and year of collection, were retrieved from NCBI. Seven-gene multi-locus sequence types (ST) were determined using Kleborate v3.2.4 [50] with the published *K. oxytoca* SC scheme [51]. Accession and QC details are available in **Supplementary Table 2**.

### Identification of *K. oxytoca* Species Complex K-loci

We used BLASTn search implemented within Kaptive v2.0.3 [34] to identify the conserved K locus flanking regions and extract the connecting assembly regions for each genome in the preliminary dataset (Kaptive used with the *K. pneumoniae* SC K locus reference database, gap fill size of 500 bp and otherwise default parameters). To remove redundant sequences, putative full-length K loci were clustered using MeShClust v3.0 [52], with a nucleotide identity and coverage threshold of 90% for determining cluster membership.

The centre sequence of each cluster was then extracted and annotated with Prokka v1.14.6 [53] using default parameters, and the *K. pneumoniae* SC K locus primary reference sequences as trusted proteins. We then performed an all-versus-all pairwise comparison of translated coding sequences (CDSs) with MMseqs2 [54]. This indicated an inflection point in the distribution at 82.5% amino acid identity, which we used as the threshold for defining distinct genes (**Supplementary Figure 2**). Predicted K-locus proteins were screened using HMMER v3.4 hmmscan against Pfam-A, PGAP and custom Wzy HMMs [55]. Putative Wzy polymerases were assigned from Wzy HMM hits.

Glycosyltransferases were identified from protein sequence clusters supported by Pfam-A and/or PGAP annotations consistent with glycosyltransferase function. Glycosyltransferase CDSs clustering with the WbaP reference PRK15204.1 were assigned as WbaP-like UDP-galactose:undecaprenyl-phosphate galactose phosphotransferases, while glycosyltransferase CDSs clustering with the WcaJ reference PRK10124.1 were assigned as WcaJ-like UDP-phosphate glucose phosphotransferases. Genomes carrying *wzy*-deficient K loci were screened for prophages using PHASTESTv3.0 [56], annotations cross-referenced to the Wzy HMM results, and a bitscore threshold of >35 [55].

### Generation of the K locus database

Consistent with the Kaptive typing framework, distinct K loci were defined in the preliminary dataset as distinct sets of genes (using the 82.5% translated amino acid identity threshold defined above). Loci were included in the database if there was at least one representative with all genes found within a single contiguous assembly region, and without predicted truncation of any of the core capsule synthesis and assembly genes, *galF, wzi, wza, wzb, wzc* and *ugd*, that would otherwise be predicted to result in a capsule null phenotype (note that we did not require that all loci contain annotated *wzx* and *wzy* because these can be difficult to identify, and in a minority of cases are located elsewhere in the genome [55, 57]). Representatives without insertion sequence (IS) insertions were preferentially selected where available. Where no IS-free version of a locus was found, we generated a synthetic IS-free locus by excision of the complete IS sequence (including the inverted repeat sequences) plus one copy of the associated direct repeat (determined using ISFinder [58]).

Each distinct K locus was labelled as KL, followed by a unique number starting at 01. K loci corresponding to those of the original *Klebsiella* serotype reference strains were assigned numbers matching the corresponding serotype identifier (i.e. KLx matches the locus of the Kx serotype reference strain). For each K locus, we generated a GenBank format file (.gbk) which included all annotated coding sequences and the full nucleotide sequence. Annotations were manually curated to standardise gene names and ensure orthologous coding sequences were annotated to the same length, with equivalent start codons. Where known, we annotated predicted capsule phenotypes within the GenBank files, which were concatenated into a multi-record file for use within Kaptive (available at github.com/klebgenomics/KoSC-surface-antigen-loci) and implemented in Kaptive Web (kaptive-web.erc.monash.edu).

Where *K. oxytoca* SC K loci were orthologous to those in the *K. pneumoniae* SC, we made a note in the GenBank file. With the exception of the loci corresponding to the serotype reference strains (described above), we chose not to enforce the use of the same KL numbers for orthologous loci because; i) the numbers of the *K. pneumoniae* SC orthologs were much greater than the total number of *K. oxytoca* SC loci; and ii) we expect that novel *K. oxytoca* SC and *K. pneumoniae* SC loci will continue to be discovered with expanding sequencing efforts and as novel loci evolve within these populations, so it will not be possible to synchronise numbers on an ongoing basis (i.e. where a corresponding number is already in use).

### Database testing

We tested the novel K locus database, which was initially developed using the preliminary dataset, by screening 4,055 high-quality *K. oxytoca* SC genomes in the final dataset using Kaptive v3.1.0 [30]. Genomes that were marked as ‘Untypeable’ were further investigated for the presence of putative novel loci by extraction of contiguous locus sequences, annotation and comparison of the gene content to existing loci. Where a sequence was confirmed to represent a distinct set of genes defined at the 82.5% translated amino acid identity threshold, its annotation was curated and added to the reference database as above (n=8 additional loci were added). For comparison, we also typed the final *K. oxytoca* SC dataset using the *K. pneumoniae* SC K locus database with *Kaptive* v3.1.0.

We report and compare typing performance using a dereplicated subset of the final dataset to reduce the impact of sampling biases in the public genome collections: e.g. where outbreak investigations and/or studies focussed on specific antimicrobial-resistant strains can lead to overrepresentation of a subset of AMR-associated clones. To dereplicate, we selected a single representative of each unique combination of genomic cluster, country of origin, year of collection, 7-gene ST and best matching K locus (regardless of the reported match confidence). Genomic clusters were defined using Mash v2.3 [59] (sketch size 10000, *k*-mer size 21, Mash threshold 0.003) and K loci were defined using the novel *K. oxytoca* SC database. Rarefaction curves showing K locus accumulation with increasing genome sampling in the dereplicated dataset ‘were calculated for each of the four most common species using the “rarecurve” function of the vegan v2.7.2 R package [60].

### Seroepidemiology analysis

To showcase the utility of our novel K locus database to support seroepidemiology analyses, we identified three diverse publicly available *K. oxytoca* SC clinical isolate whole genome sequence collections from distinct geographies: Australia (n=92) [2], China (n=102) [61], and Italy (n=61) [3] (**Supplementary Table 3**). We typed all three genome collections using our novel *K. oxytoca* SC database with *Kaptive* v3.1.0, calculated and compared the relative prevalences of each K locus overall and by geography.

### Data visualisation and statistical analyses

Data were summarised and visualised, and statistical analyses performed using R v4.5.1 [62] with the following packages: tidyverse v2.0.0 [63], reshape2 v1.4.4 (https://github.com/hadley/reshape), vegan v2.7.2 [60] and patchwork v1.3.2 (https://patchwork.data-imaginist.com). Locus gene cluster and synteny plots were generated with clinker v0.0.32 [64].

## RESULTS

### *K. oxytoca* Species Complex K locus database comprises 88 distinct loci

A total of 88 distinct K loci were identified, ranging in length from 21,171 bp (KL37) to 30,521 bp (KL40) (median 25,901 bp) and containing between 17 (KL37 and KL11) and 25 (KL31) ORFs (median 21). All loci contained the conserved capsule synthesis and export machinery genes *galF, cpsACP, wza, wzb, wzc, gnd and ugd*). The *wzi* gene, implicated in capsule attachment to the cell surface, was present in 87 loci (98.9%) and absent from KL81, which instead included four ORFs encoding proteins implicated in exopolysaccharide and/or group 4 capsule production in *Escherichia coli* [65, 66] (**Figure 1A**). Orthologs of the same genes (97% nucleotide identity) were previously described within *K. pneumoniae* SC KL33, which also lacks *wzi* but nonetheless results in production of the K33 capsule [31]. *wzx* and *wzy* are notoriously difficult to annotate due to high sequence divergence [55] and were detected in only 86 (97.7%) and 82 (93.1%) loci, respectively. Loci lacking *wzy* were included in the database as it has been shown in *Acinetobater baumannii* that functional copies of *wzy* can be located on mobile elements outside of the K locus, such as on prophages [57, 67]. In fact, upon further inspection, several genomes carrying *wzy*-deficient loci were found to have a copy of *wzy* located on a mobile element (e.g. SAMEA4781318; KL28, see Figure 4, Supplementary Material).

**Figure 1:**
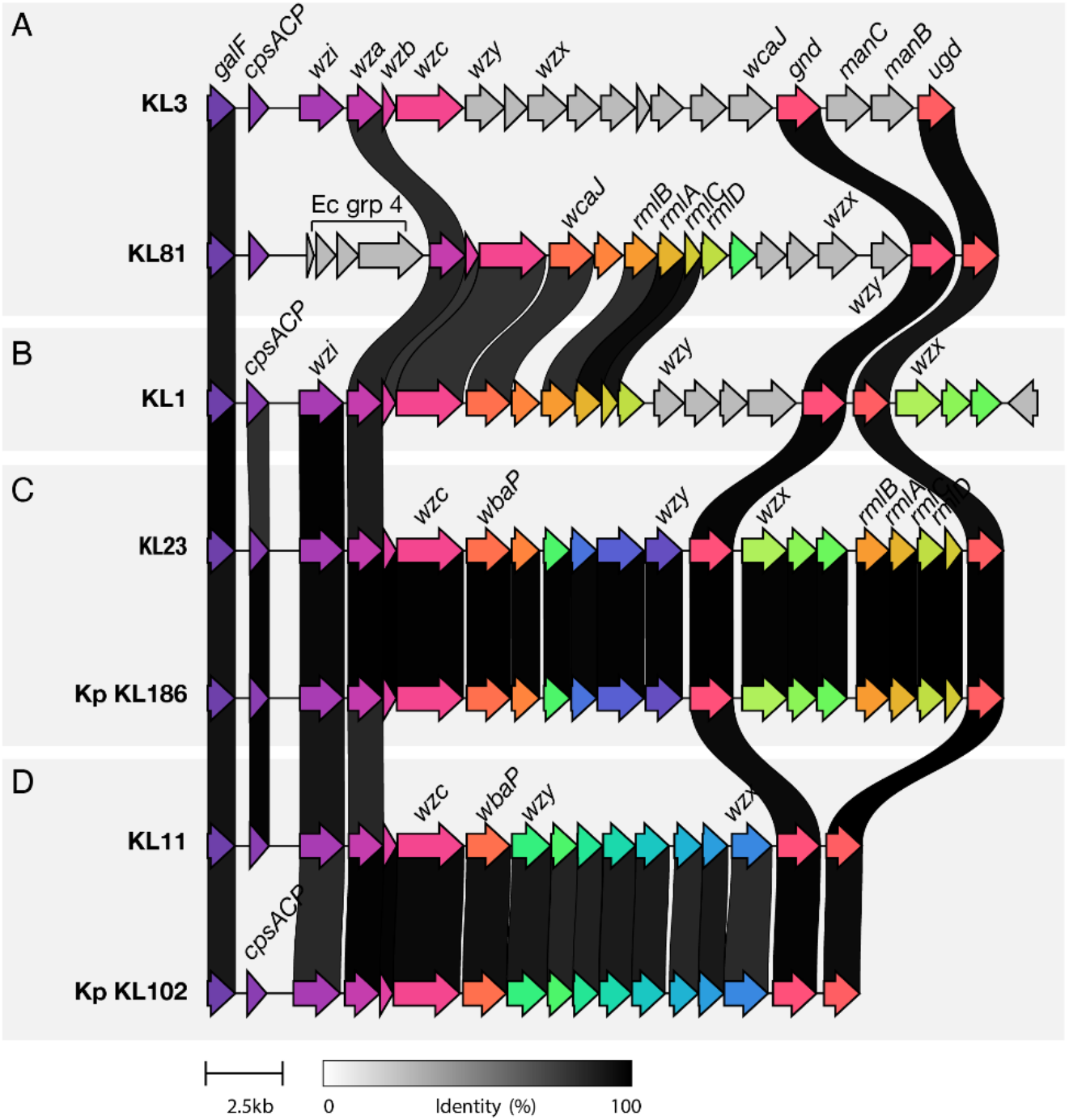
Klebsiella oxytoca Species Complex (K. oxytoca SC) K loci examples and comparisons to K. pneumoniae SC K loci. Arrows represent genes coloured by orthology groups as defined by clinker [64]. Labels indicate genes encoding conserved capsule synthesis and assembly machinery and/or genes discussed in the text (remaining genes not labelled for legibility). Black lines connect orthologous genes sharing amino acid identity, as indicated in the legend (note only those gene pairs sharing ≥82.5% identity are shown). **A)** A typical K. oxytoca Species Complex (SC) K locus (KL3) and the atypical KL81, which lacks wzi and instead contains four genes with orthology to those implicated in Escherichia coli group 4 capsule synthesis (labelled Ec grp 4). **B)** An example K. oxytoca SC with several genes located downstream of ugd. **C)** An example of K. oxytoca SC and K. pneumoniae (Kp) SC direct orthologs (i.e. a pair of loci for which all genes are shared above the 82.5% translated amino acid identity threshold). **D)** K. oxytoca SC KL11 is a distant ortholog of K. pneumoniae (Kp) SC KL102, with predicted functional orthology: the loci differ only by sequence divergence in cpsACP (<82.5% translated amino acid identity) which is a core K locus gene.

Initiating glycosyltransferases were identified in all loci; n=77 *wbaP* and n=11 *wcaJ*, indicating a dominance of galactose initiating monosaccharides over glucose. Each locus also contained between one and five additional putative glycosyltransferases from a total pool of 288 distinct putative glycosyltransferase genes (82.5% amino acid identity threshold). Individual glycosyltransferases were found in ≤7 K loci each. The *rmlACBD* genes (associated with rhamnose synthesis) were identified in 57 loci (64.8%), while the *manCB* genes (associated with mannose synthesis) were identified in 34 loci (38.6%, of which 14 also contained *rmlACBD*).

We observed ten K loci which had putative capsule biosynthesis genes located downstream of *ugd* (KL1, KL16, KL21, KL22, KL24, KL40, KL44, KL63, KL68, KL82, see example in **Figure 1B**). These included the recently described K locus corresponding to *K. grimontii* strain K15g [28] (designated here as KL1 and annotated with the corresponding published structural phenotype, labelled *K. oxytoca* SC K1). Others have previously described *K. pneumoniae* K loci with genes downstream of *ugd* [68], and together with our findings highlight the variable nature of these biosynthetic loci, which are thought to be subject to reassortment and rearrangement through chromosomal recombination and IS-mediated transposition [32, 69].

There were 10 *K. oxytoca* SC K loci that were completely orthologous to those in the *K. pneumoniae* database (**Supplementary Table 4** and see example in **Figure 1C**), here defined using the logic of the Kaptive typing framework which states that two loci are the same if they comprise the same set of genes at a defined translated amino acid identity threshold (82.% for *Klebsiella* spp.). The orthologous loci include those for which the original *Klebsiella* serotype reference strains are now recognised as members of the *K. oxytoca* SC (labelled KL26, KL29, KL41, KL66, KL70, KL74 in both databases) as well as *K. oxytoca* SC KL3, KL10, KL19 and KL23 which are orthologous to *K. pneumoniae* SC KL174, KL152, KL181 and KL186, respectively. We expect that these orthologous pairs of loci result in the production of polysaccharides with matching structures; however, to date structures have been described only for those corresponding to KL26, KL41, KL66, KL70 and KL74 and hence only these loci are annotated with corresponding K type predictions in the *Kaptive* database.

A further seven *K. oxytoca* SC loci were distant orthologs to those from the *K. pneumoniae* SC database, which we define as loci sharing a full set of gene orthologs but with ≥1 below the 82.5% amino acid identity threshold (**Supplementary Table 4** and example in **Figure 1D**). These included *K. oxytoca* SC KL6, KL11 and KL14, which differed from their *K. pneumoniae* SC locus orthologs only by sequence divergence in the core K locus gene, *cpsACP*, for which we expect functional equivalence (although the precise role of this gene in capsule biosynthesis remains unknown [33]). We, therefore, predict that these *K. oxytoca* SC loci result in the production of orthologous polysaccharide structures to those encoded by *K. pneumoniae* SC KL43, KL102 and KL109, respectively. Polysaccharide structures have been defined for *K. pneumoniae* SC strains carrying KL43 and KL102; hence we have annotated the predicted structures for the *K. oxytoca* SC orthologs in the *K. oxytoca* SC database. The remaining distant ortholog pairs differed by sequence divergence in ≥1 capsule-specific synthesis gene and including ≥1 glycosyltransferase gene, for which functional equivalence is difficult to predict (**Supplementary Table 4**). Additionally, we determined that *K. oxytoca* SC KL4 was similar to *K. pneumoniae* SC KL145; however, *K. pneumoniae* SC KL145 appears to carry truncated versions of three glycosyltransferase genes. We, therefore, consider *K. oxytoca* SC KL4 and *K. pneumoniae* SC KL145 as an additional distant orthologous pair, but which likely encode distinct polysaccharides.

### *K. oxytoca* SC K locus database enables comprehensive typing of *K. oxytoca* SC genomes

Among 4,055 genomes in the final *K. oxytoca* SC dataset that passed quality control, 3,981 (98.1%) were successfully typed to 88 distinct K loci using the *K. oxytoca* SC database (Kaptive confidence = ‘Typeable’). In the dereplicated set of 2,244 genomes, which comprised 10 *K. oxytoca* SC taxa (plus *K. indica*) and 670 7-gene STs, representing isolates collected from ≤1886 to ≥2024 and from ≥65 countries, n=2,192 (97.6%) were successfully typed (**Supplementary Table 2**). As expected, the typing rate achieved using the *K. oxytoca* SC database was significantly higher than that achieved using the *K. pneumoniae* SC database; 97.6% vs 50.3% typed from the dereplicated *K. oxytoca* SC dataset (p-value < 0.0001 by Fisher’s Exact test, OR 41.6, 95% confidence interval (CI) 31.14–56.59). The total number of loci reported was also substantially higher when using the *K. oxytoca* SC database compared to *K. pneumoniae* SC database (88 vs 22, respectively). Furthermore, matches to the *K. oxytoca* SC database were generally higher quality than those to the *K. pneumoniae* SC database (**Supplementary Results**), providing further evidence of the benefit of using a tailored organism-specific database.

To further estimate the population coverage of the *K. oxytoca* SC K locus database we generated rarefaction curves showing the rate of accumulation of distinct K loci with the addition of novel genomes for the four most-sampled species in the dereplicated set (**Figure 2**); *K. michiganensis* (n=1,088 genomes), *K. oxytoca* (n=689), *K. grimontii* (n=348) and *K. pasteurii* (n=81) (n≤15 for each of the other taxa). The curve for *K. oxytoca* appears to have reached a plateau at n=36 K loci, while those for *K. michiganensis* and *K. grimontii* are trending towards plateau (at much higher levels of >40 and >50 K loci, respectively), suggesting that the current database captures most K loci present among the available genomes.

**Figure 2:**
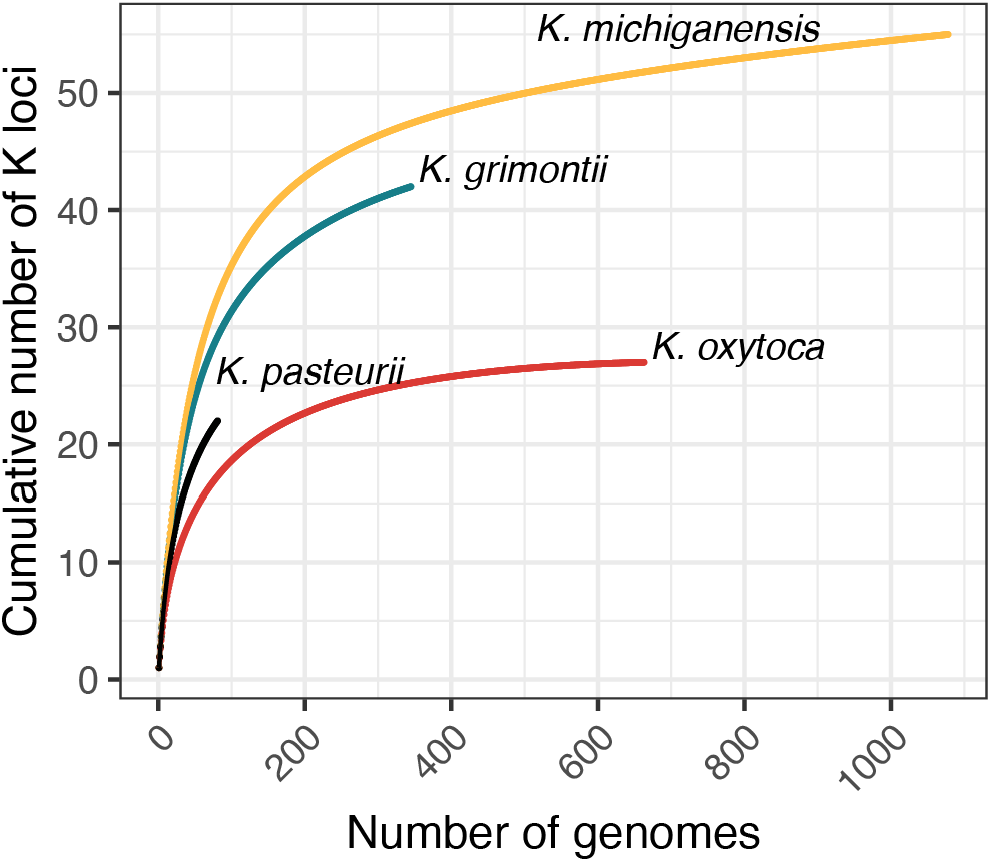
Rarefaction curves showing accumulation of K loci with increasing sampling of Klebsiella oxytoca Species Complex genomes. The four most common species among the available genome data are represented independently as indicated by the labels. The curve for K. oxytoca has reached a plateau. The curves for K. michiganensis and K. grimontii are trending to, but do not reach, plateau. The curve for K. pasteurii has not reached plateau, likely due to its comparatively small sample size.

### K loci frequency and species distribution

The *K. oxytoca* SC K locus frequency distribution was negatively skewed; most loci (n=54, 61%) were observed in <1% typeable genomes in the dereplicated dataset, 20 loci (23%) were observed in ≥1% but <2% genomes, and only 14 loci (16%) were observed in >2% genomes (**Figure 3**). Nevertheless, half of all K loci were represented among ≥2 species each (n=44 of 88 K loci), with 24 K loci (28%) each identified in ≥3 species. Notable exceptions were KL34 and KL69, which were present in 5% and 3% of genomes, respectively, all of which were *K. oxytoca sensu stricto*. However, it is unclear if this observation represents a genuine species restriction or an artefact of our convenience sample.

**Figure 3:**
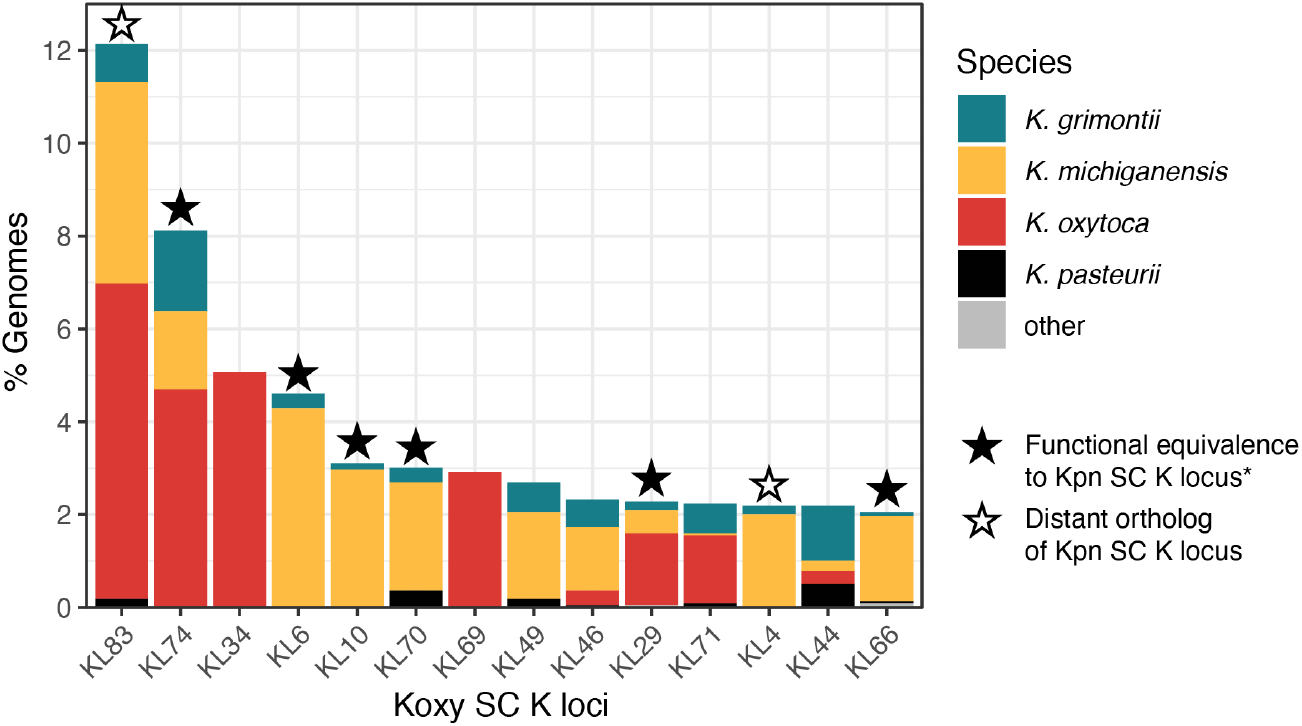
Relative frequency and species representation of the top 14 Klebsiella oxytoca Species Complex (Koxy SC) K loci. K loci observed in >2% of typeable genomes in the dereplicated dataset are shown (total n=2,192 typeable genomes). Koxy SC species representation is indicated by colour as per the legend. Stars indicate Koxy SC K loci with orthology to those defined for the Klebsiella pneumoniae Species Complex (Kpn SC); filled stars indicate direct orthologs (share an identical set of genes at the 82.5% amino acid identity threshold) or those likely to be functionally equivalent (all genes shared with ≥82.5% amino acid identity except the core gene cpsACP, which had <82.5% identity); open stars indicate more distant orthologs for which >1 gene was shared with <82.5% amino acid identity. * Functional equivalence is known for KL74, KL70, KL29 and KL66, and predicted for KL6 and KL10.

Many of the most common K loci were orthologs of those defined for the *K. pneumoniae* SC (six direct orthologs and/or likely functionally equivalent loci; two distant orthologs among the top 14 K loci, **Figure 3**). In fact, 31% of all typeable genomes in the dereplicated dataset (n=674) harboured a direct ortholog and/or predicted functionally equivalent K locus to one of those defined among the *K. pneumoniae* SC, and 16% (n=346) harboured distant orthologs (as defined above).

### Use case: seroepidemiology of clinical *K. oxytoca* SC

To showcase the utility of our database we explored capsule epidemiology among three clinical isolate collections where >60 diverse *K. oxytoca* SC isolates were sequenced: n=102 from China, representing clinical isolates collected between January 2020 and November 2021 from a study spanning 69 hospitals [61]; n=92 from Melbourne, Australia representing all third-generation cephalosporin-resistant isolates (n=22), plus all blood infection isolates not already included (n=18), and additionally randomly selected clinical isolates from a single hospital network in 2016 [2]; and n=61 clinical isolates from a single hospital in Pavia, Italy, representing all clinical isolates collected between June 2017 and November 2018 via standard diagnostics protocols [3] (**Supplementary Table 3**). All four of the common species were represented within each dataset, with *K. michiganensis* and *K. oxytoca* together accounting for 82–89% of genomes. The genomes represented 150 distinct 7-gene STs, with 44–71 distinct STs per dataset. All 255 genomes had been included in the final dataset used to develop the database.

All 255 genomes were successfully typed with the *K. oxytoca* SC K locus database, indicating a high diversity of loci (23–38 distinct loci per study) and capturing 54 (61%) of the total 88 loci in the *K. oxytoca* SC database. While individual K locus frequencies varied by site, there was overlap among the sets of most common loci (**Figure 4A**). KL74 and KL34 were the two most common loci in both the Australian (n=12 and n=8 of 92 genomes, respectively) and Chinese datasets (n=11 each of 102 genomes), with KL74 also ranking joint third in the Italian set (n=4 of 61). KL83 was the most common locus in the Italian set (n=12) and ranked joint fourth in the Australian set (n=5), and third in the Chinese set (n=8). Together KL74, KL34 and KL83 accounted for 28.2% isolates from the three clinical collections and 25.2% (n=736 of 2,919) of typeable genomes in our larger dereplicated dataset. Each of these three common loci were associated with multiple 7-gene STs but the ST compositions differed by dataset (**Figure 4B**), consistent with differences in locally circulating clones (although the sample sizes were too small to support statistical comparisons, n≤8 genomes per ST per dataset).

**Figure 4:**
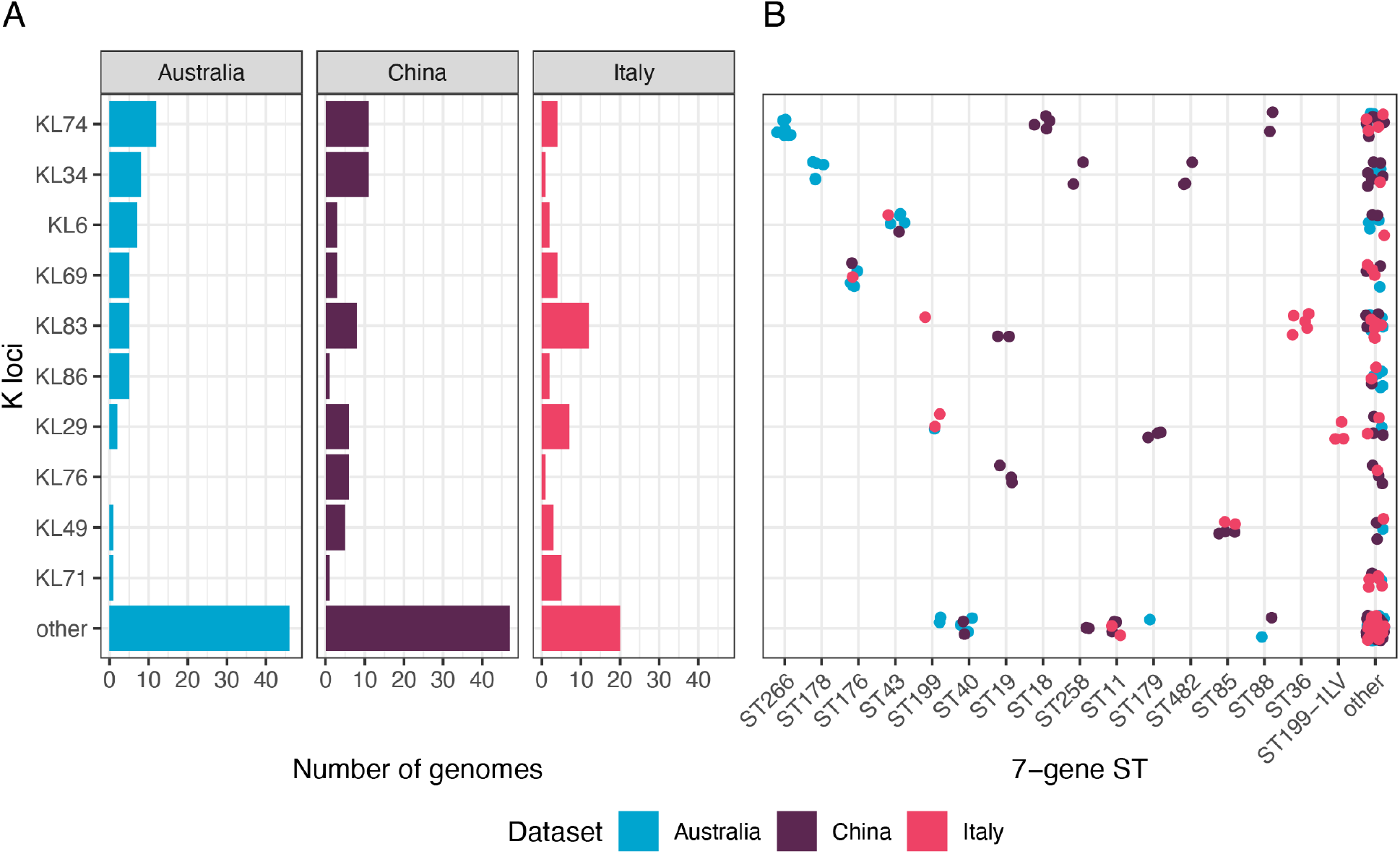
K locus frequencies among clinical Klebsiella oxytoca Species Complex isolate collections and associations with 7-gene multi-locus sequence types. Isolate genomes were sourced from three published studies from Australia (n=92) [2], China (n=102) [61] and Italy (n=61) [3]. **A)** Frequencies of K loci represented by ≥5 genomes in at least one dataset (remaining loci grouped as ‘other’). **B)** Sequence types (STs) associated with each K locus, coloured by dataset as indicated in legend. STs represented by ≥3 genomes in at least one dataset are individually labelled (remaining STs grouped as ‘other’).

## DISCUSSION

We have presented a novel *K. oxytoca* SC-specific Kaptive-compatible K locus database that facilitates comprehensive typing of *K. oxytoca* SC genomes. This novel database contains 88 distinct *K. oxytoca* SC K loci defined as unique sets of genes at the 82.5% amino acid identity threshold. Consistent with current evidence from *K. pneumoniae* SC [31], we anticipate that the majority of these loci result in the production of distinct capsule polysaccharides. Seven *K. oxytoca* SC capsule polysaccharide structures have been described to date and three additional structures are predicted here based on orthology between the associated *K. oxytoca* SC K loci and those of *K. pneumoniae* SC isolates expressing defined structures (**Figure 1, Supplementary Table 4**). All known and predicted structures are annotated within the *K. oxytoca* SC K locus database and are reported by Kaptive in the ‘Best match type’ column.

Our data suggest that the *K. oxytoca* SC-specific database captures the majority of K loci present among currently available *K. oxytoca* SC genome sequences (98% genomes successfully typed; **Figure 2**). Public genome collections are known to be biased towards human-derived isolates (70% of dereplicated genomes in this dataset for which source data were available) and those collected from high-resource settings, and we anticipate that novel *K. oxytoca* SC K loci will be identified with greater diversification of the underlying genome collections. We plan to perform periodic screens for novel *K. oxytoca* SC loci as we have done for *K. pneumoniae* SC [34], and encourage database users to share candidate novel loci for curation and inclusion in the *K. oxytoca* SC database (details provided at https://github.com/klebgenomics/KoSC-surface-antigen-loci).

Of note, our analyses showed that when typing *K. oxytoca* SC genomes, the novel *K. oxytoca* SC-specific K locus database results in almost twice as many typed genomes with superior quality matches than the previously published *Klebsiella* database (which was primarily designed for *K. pneumoniae* SC and has now been explicitly relabelled as the *K. pneumoniae* SC database to avoid confusion). Furthermore, inspection of the Kaptive results suggests that the *K. pneumoniae* SC database can result in mistyping when applied to *K. oxytoca* SC genomes. Kaptive uses the proportion and counts of expected and ‘extra’ genes as part of its scoring algorithm, searching only for those genes present in the reference database. We have previously shown that the default Kaptive settings result in high sensitivity and accuracy (≥98% for draft genomes of standard quality) for databases with high population coverage: i.e. the *K. pneumoniae* SC and *Acinetobacter baumanii* databases applied to *K. pneumoniae* SC and *A. baumanii* genomes, respectively [30]. In that systematic testing, genomes carrying loci that were represented in the reference databases were very rarely mistyped, but a small number of novel loci were mistyped. We, therefore, strongly caution against using databases to type species for which they were not designed and/or where a large number of novel loci or novel polysaccharide synthesis genes are expected, without further optimisation of Kaptive’s settings and manual inspection of the outputs.

Our dereplicated dataset represents a convenience sample of *K. oxytoca* SC genomes from which individual K locus frequencies should be interpreted with caution as they may not be indicative of the true frequencies in *K. oxytoca* SC populations from broad geographies and niches. Nonetheless, the data clearly show that *K. oxytoca* SC harbour a high diversity of K loci, with capacity for horizontal transfer between species (**Figure 3**).

Almost one third of *K. oxytoca* SC genomes (31%) harboured K loci predicted to result in production of polysaccharide structures matching those encoded by loci that have been previously associated with *K. pneumoniae* SC. Notably, *K. oxytoca* SC KL11 (1.8% *K. oxytoca* SC genomes) was predicted functionally equivalent to *K. pneumoniae* SC KL102, associated with the globally distributed multi-drug resistant ST307 clone [70] and among the top ranked K loci from neonatal sepsis isolates in Africa and South Asia [71]. In contrast, the remaining functionally equivalent K loci are rarely reported among clinical *K. pneumoniae* SC isolate collections [35, 36, 71–73].

In order to further demonstrate the utility of the *K. oxytoca* SC K locus database to support seroepidemiology analyses, we used it to estimate the relative frequencies of K loci among *K. oxytoca* SC causing human infections from three published datasets (**Figure 4**). These data suggest that there is substantial capsule type diversity, with variation between study locations that may in part be driven by differences in locally circulating clones, as is also the case for *K. pneumoniae* [71, 72]. However, these analyses were based on genome collections of small size and representing diverse sampling strategies, and as such should be considered illustrative rather than representative of the broader *K. oxytoca* SC population. Larger systematic genome collections will be required to understand the true population K locus diversity and power robust frequency estimates: e.g. using outbreak-corrected Bayesian frequency estimates as recently described for *K. pneumoniae* [71]. The novel *K. oxytoca* SC database presented here will facilitate and support such analyses, which will be essential to understand capsule-specific virulence potential and ecology, and to prioritise capsule types for inclusion in vaccines or as targets for therapeutic strategies.

## AUTHOR STATEMENTS

### Author contributions

Conceptualisation: KEH, KLW

Data curation: MMA, NM, LH

Formal analysis: MMA, NM, TDS, KLW

Funding acquisition: KEH, KLW

Methodology: MMA, NM, TDS, KLW

Project administration: KLW

Validation: NM, KLW

Visualisation: NM, KLW

Writing - original draft: MMA, KLW

Writing - revising and editing: All authors

