## Supplementary material for "A capsule polysaccharide synthesis locus database for the *Klebsiella oxytoca* Species Complex": 20261007_Supplementary_Materials.pdf

**Repositories:** The novel capsule (K) locus database is available at github.com/klebgenomics/KoSC-surface-antigen-loci

### SUPPLEMENTARY RESULTS

#### Organism-specific database supports superior K locus typing

Among those *K. oxytoca* SC genomes reported as typeable from the dereplicated dataset, matches were generally higher quality when using the *K. oxytoca* SC database than when using the *K. pneumoniae* SC database: more than half, (n=1,177, 53.7%) of the loci typeable with the *K. oxytoca* SC database were very high-quality matches (i.e. the loci were found in a contiguous piece without any missing genes nor any truncated nor extra genes compared to the best matching reference locus, as indicated by no entry in the 'Problems' column of the Kaptive output). In contrast, using the same criteria, only 176 (15.59%) of the loci typeable with the *K. pneumoniae* SC database were very high-quality matches. Additionally, the weighted locus coverages and identities were significantly higher for typeable matches to the *K. oxytoca* SC database than to the *K. pneumoniae* SC database (median coverage 100% vs 96.81%, p-value < 0.001; median identity 99.78% vs 96.97%, p-value < 0.0001 by Wilcoxon Ranked Sums tests).

Interestingly, we identified 300 genomes for which results generated using the *K. oxytoca* SC database (typeable or untypeable) were discordant to those reported using the *K. pneumoniae* SC database (typeable only, to exclude the high number of low confidence matches reported with this database): i.e. where the best matching K locus reported using the *K. pneumoniae* SC database was neither a direct nor a distant ortholog of the corresponding locus reported using the *K. oxytoca* SC database (locus orthologs defined as per main text), suggesting that one of the results was incorrect. We explored the quality of these matches: 147 *K. oxytoca* SC K locus matches (49.0%) were reported with very high quality (as defined above), whereas all of the 300 corresponding *K. pneumoniae* SC K locus matches were reported with one or more problems, including 296 matches missing one or more genes from the best matching reference locus and 228 matches that included one or more additional genes from other reference loci.

More broadly, for 271 of the 300 (90.3%) discordant matches between the databases, the weighted coverage of the best match reference locus genes was greater, and the absolute number of other locus genes was equal or fewer for the *K. oxytoca* SC database match compared to the *K. pneumoniae* SC database match (**Supplementary Figure 3**). Of the remaining 29 matches, six contained full length contiguous loci, whereas 23 contained fragmented loci which are notoriously difficult to type and therefore not investigated further (between 2 and 8 pieces each). For five of the six continuous loci, inspection of the annotated regions of the genomes supported the match to the *K. oxytoca* SC database (GCF_036668975, SAMEA3357557, SAMEA4781093, SAMEA5684288, SAMEA10468609). The sixth genome (GCF_036441055) was reported as a typable match to *K. pneumoniae* SC KL122 and was reported as untypeable (best match KL3) using the *K. oxytoca* SC database. Further investigation of the Kaptive outputs and annotated assembly region suggested that GCF_036441055 carries an ortholog of *K. pneumoniae* SC KL122. This locus was not added to the *K. oxytoca* SC reference database because the core *cpsACP* and *wzi* genes are truncated in the GCF_036441055 assembly (i.e. the locus is predicted to be nonfunctional) and no other full-length *K. oxytoca* SC orthologs were identified.

**SUPPLEMENTRAY FIGURES**

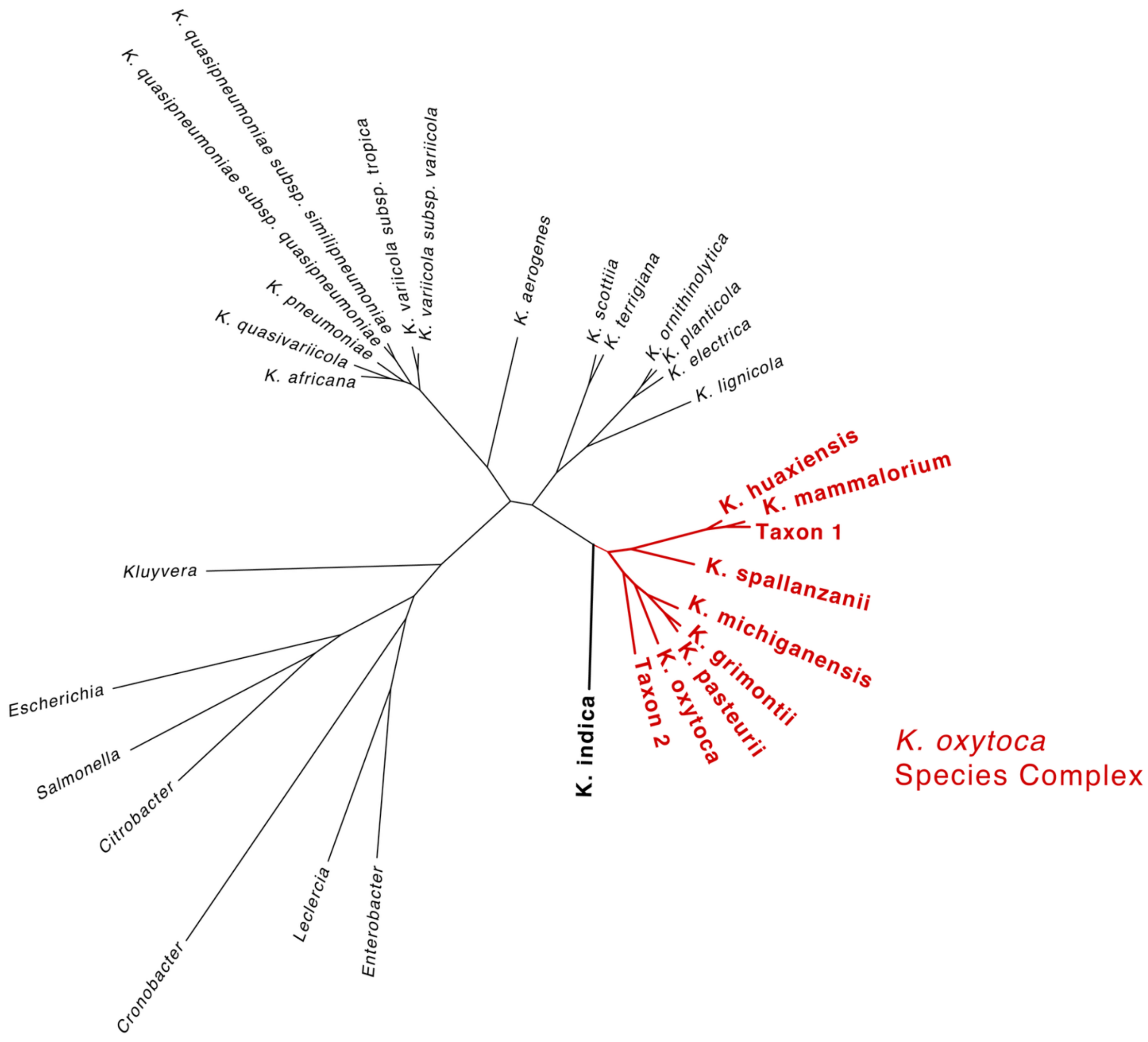


**Figure S1: Phylogenetic relationships among *Klebsiella* (K.) species.** Unrooted maximum likelihood phylogeny showing all known members of the *Klebsiella oxytoca* Species Complex (*K. oxytoca* SC, branches shown in red) and their relationships to other *Klebsiella* spp. The *K. oxytoca* SC K locus database also includes loci derived from genomes of the closely related, *Klebsiella indica* (shown in black, bold). Note: formal species description for "*Klebsiella mammalorium*" is in progress (KLW and LH).

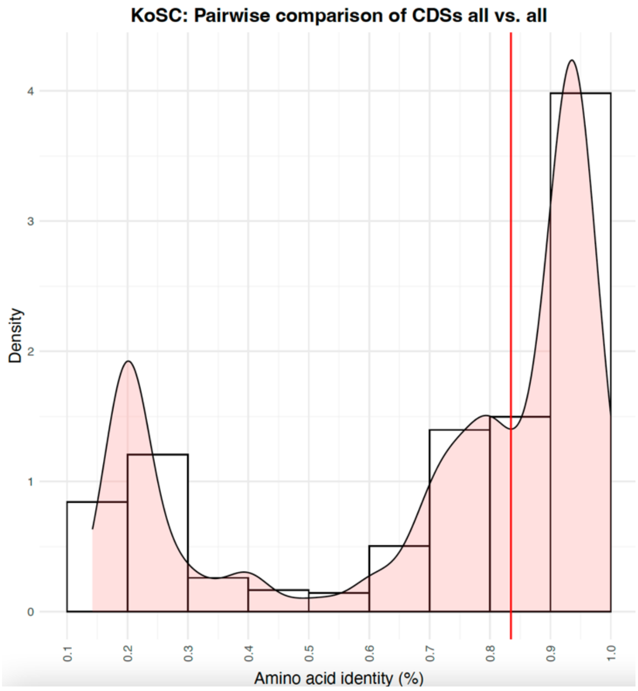


**Figure S2: Distribution of pairwise amino acid identities between *Klebsiella oxytoca* Species Complex (KoSC) K locus genes.** Coding sequences (CDS) were extracted from representative sequences of each MeshClust [1] cluster and translated identities calculated using MMSeqs2 [2]. Red vertical line indicates the amino acid identity threshold applied to distinguish unique genes.

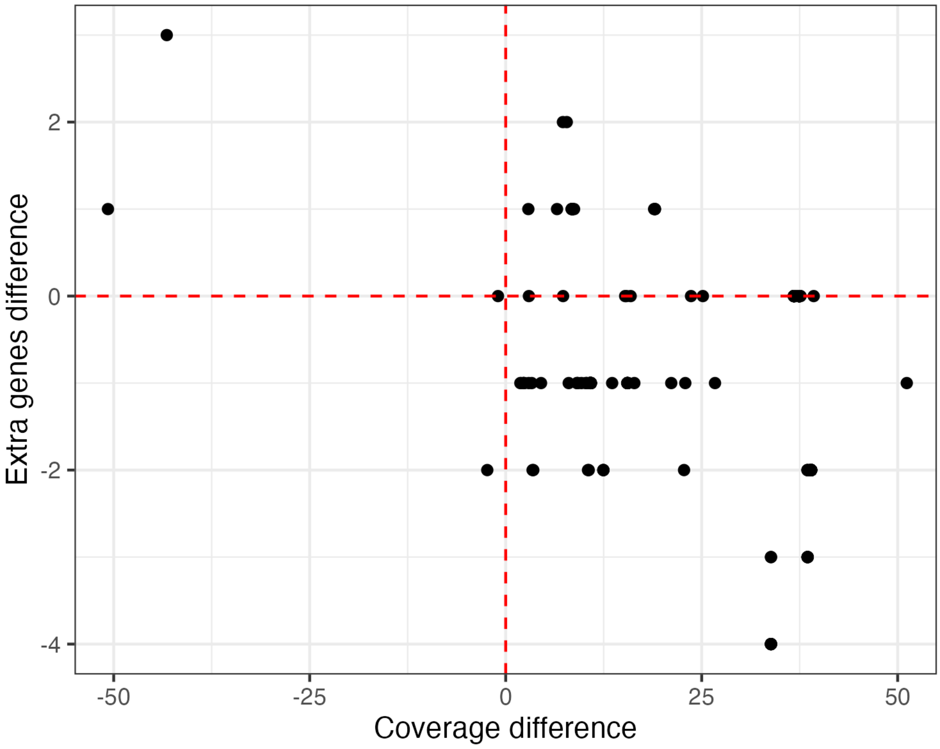


**Figure S3: Comparison of locus coverage and number of extra genes reported for genomes typed with the *Klebsiella oxytoca* Species Complex (SC) and *Klebsiella pneumoniae* SC K locus databases.** The plot shows the difference in the reported locus coverages (*K. oxytoca* SC coverage – *K. pneumoniae* SC coverage) and counts of numbers of extra genes (*K. oxytoca* SC – *K. pneumoniae* SC) for discordant typing results (i.e. where the best match loci reported using the *K. oxytoca* SC database was not a direct ortholog of that reported using the *K. pneumoniae* SC database). 'Extra genes' reflect matches to genes from a reference locus other than the reported best match locus, found within the locus region of the input assembly. The red dashed lines indicate the y = 0 and x = 0 lines. The majority of data points fall in the lower right quadrant, indicating that the *K. oxytoca* SC match was reported with higher coverage and fewer extra genes than the *K. pneumoniae* SC match.

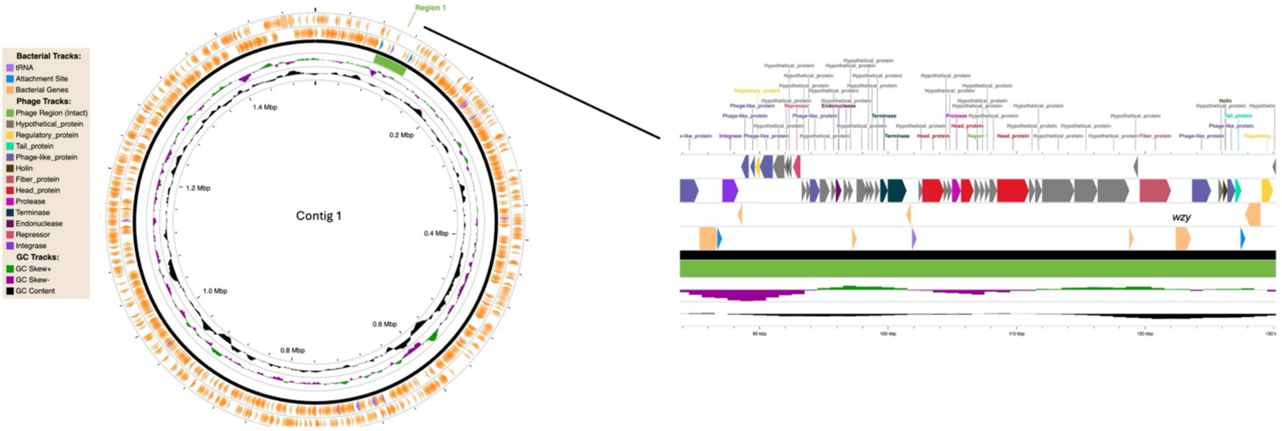


**Figure S4: Detection of *wzy* located on prophage.** This figure depicts the location of a *wzy* ortholog (detected using custom Wzy HMMS [3] on a prophage in SAMEA4781318, as determined by PHASTEST v3.0 [4]. The WzyB ortholog was confirmed via HMMER v3.4 hmmscan on contig 1 between 525012-526184 bp, with a bit-score of 79.6.

### SUPPLEMENTRAY TABLES

**Supplementary Table 1** (separate .xlsx file): Genomes included in the preliminary dataset.
**Supplementary Table 2** (separate .xlsx file): Genomes included in the final dataset, plus isolate metadata, 7-gene sequence types, Kaptive typing information for matches to the *Klebsiella oyxtoca* Species Complex (SC) and *Klebsiella pneumoniae* SC K locus databases.
**Supplementary Table 3** (separate .xlsx file): Genomes included in the seroepidemiology examples use case, plus 7-gene sequence types and K loci.

**Supplementary Table 4:** Pairs of orthologous loci among the *Klebsiella oyxtoca* Species Complex (*Kox* SC) and *Klebsiella pneumoniae* (Kpn SC) K locus databases.

| Kox SC locus | Kpn SC locus | Divergent genes[a] | Polysaccharide structure[b] |
|---|---|---|---|
| Direct orthologs of original *Klebsiella* sp. serotype reference strain loci | | | |
| KL26 | KL26 | - | ->3)-β-D-Galp-(1->2)-[4,6-O-Pyr-β-D-Galp-(1->4)-β-D-Glcp-(1->6)-α-D-Glcp-(1->4)]-α-D-GlcpA-(1->3)-α-D-Manp-(1->2)-α-D-Manp-(1-> |
| KL29 | KL29 | - | unknown |
| KL41 | KL41 | - | ->6)-α-D-Glcp-(1->3)-α-L-Rhap-(1->3)-α-D-Galp-(1->2)-[β-D-Glcp-(1->6)-α-D-Glcp-(1->4)-β-D-GlcpA-(1->3)]-β-D-Galf-(1-> |
| KL66 | KL66 | - | ->3)-α-D-Manp-(1->3)-α-D-Galp-(1->2)-[4-O-Lac-β-D-Glcp-(1->3)]-α-D-GlcpA-(1->3)-α-D-Manp-(1-> |
| KL70 | KL70 | - | ->4)-β-D-GlcpA-(1->4)-α-L-Rhap-(1->2)-α-L-Rhap-(1->2)-α-D-Glcp-(1->3)-β-D-Galp-(1->2)-α-L-Rhap-(1-> |
| KL74 | KL74 | - | ->3)-β-D-Galp-(1->2)-[4,6-O-Pyr-β-D-Galp-(1->4)-α-D-GlcpA-(1->3)]-α-D-Manp-(1->2)-α-D-Manp-(1-> |
| Other direct orthologs | | | |
| KL3 | KL174 | - | unknown |
| KL10 | KL152 | - | unknown |
| KL19 | KL181 | - | unknown |
| KL23 | KL186 | - | unknown |
| Distant orthologs (predicted functional equivalence) | | | |
| KL6 | KL43 | *cpsACP* | ->3)-α-D-Galp-(1->3)-[β-D-Manp-(1->4)-β-D-GlcpA-(1->2)]-α-D-Manp-(1->2)-α-D-Manp-(1-> |
| KL11 | KL102 | *cpsACP* | ->4)-[α-D-Glcp-(1->4)-β-D-GlcA-(1->3)][β-D-Glcp-(1->6)]-α-D-Galp-(1->6)-β-D-Galp-(1->3)-β-D-Galp-(1-> |
| KL14 | KL109 | *cpsACP* | Unknown |
| Distant orthologs (functional equivalence unknown) | | | |
| KL30 | KL47 | *cpsACP*, hypth, gtr | Unknown |

| | | | |
|---|---|---|---|
| KL57 | KL164 | *wzy, wcuF, wcqJ, wcuT, wcuC,* hypth | Unknown |
| KL83 | KL68 | *cpsACP, mshA* | Unknown |
| KL9 | KL26 | *wctD* | Unknown |
| Distant orthologs (functional equivalence unlikely) | | | |
| KL4 | KL145 | gtr truncations | Unknown |

[a] Genes that share ≤82.5% translated amino acid identity. Hypth = hypothetical protein, gtr = unnamed glycosyltransferase.
[b] Polysaccharide structures reported in [5–10].
